# Sprouting and anastomosis in the Drosophila trachea and the vertebrate vasculature: similarities and differences in cell behaviour

**DOI:** 10.1101/458281

**Authors:** Maria Paraskevi Kotini, Maarja Andaloussi Mäe, Heinz-Georg Belting, Christer Betsholtz, Markus Affolter

**Author notes:** Correspondence to MA and CB. MPK and MA-M contributed equally.

## Abstract

Branching morphogenesis is a fascinating process whereby a simple network of biological tubes increases its complexity by adding new branches to existing ones, generating an enlarged structure of interconnected tubes. Branching morphogenesis has been studied extensively in animals and much has been learned about the regulation of branching at the cellular and molecular level. Here, we discuss studies of the Drosophila trachea and of the vertebrate vasculature, which have revealed how new branches are formed and connect (anastomose), leading to the establishment of complex tubular networks. We briefly describe the cell behaviour underlying tracheal and vascular branching. Although similar at many levels, the branching and anastomosis processes characterized thus far show a number of differences in cell behaviour, resulting in somewhat different tube architectures in these two organs. We describe the similarities and the differences and discuss them in the context of their possible developmental significance. We finish by highlighting some old and new data, which suggest that live imaging of the development of capillary beds in adult animals might reveal yet unexplored endothelial behaviour of endothelial cells.

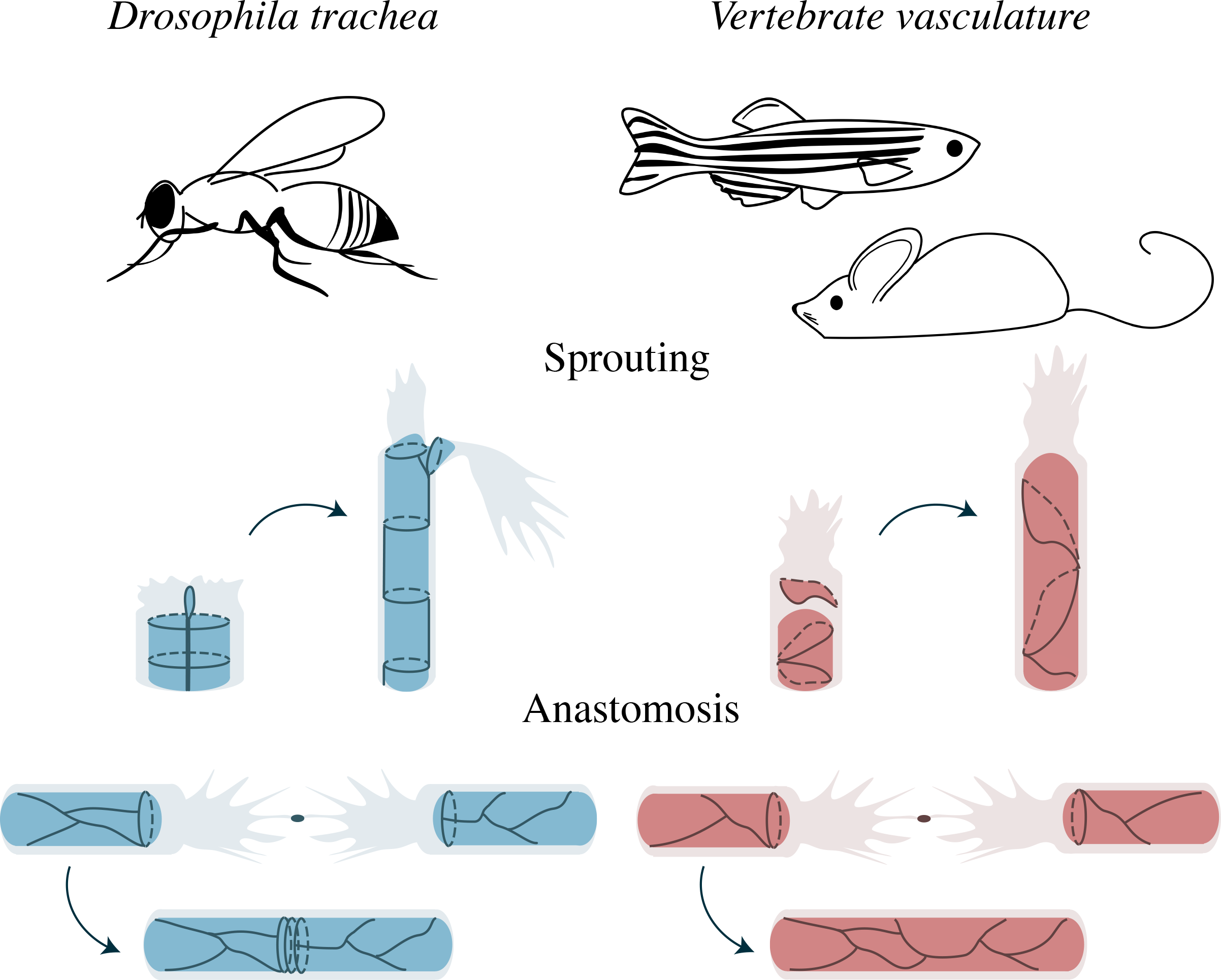

## 1 Introduction

The existence of branched biological structures, both in the plant kingdom as best visible in trees, and in the animal kingdom as exemplified by internal organs of multicellular animals such as the trachea in insects or the lung and the vasculature in mammals, has fascinated scientist for many centuries. In sharp contrast to this longstanding interest, a better understanding of how such branched structures are formed during development has only become possible in recent years. This insight was gained due to the establishment of genetic, molecular and live imaging techniques allowing for a detailed dissection of the underlying biological processes. In the past three decades, much has been learned about the control of the branching processes underlying some of these beautiful structures in plants and trees (Atkinson et al., 2014; Hollender and Dardick, 2015; Morris et al., 2017; Teichmann and Muhr, 2015) and in animals (Costantini and Kopan, 2010; Herriges and Morrisey, 2014; Morrisey and Hogan, 2010).

In this short review article, we discuss and compare the formation of two branched organs, the developing tracheal system in the drosophila embryo and the developing vasculature in vertebrates (Fig. 1). These two branched structures share many common features, including that their branching programme is to a large extent driven by very few cells in rather simple organisations; both branching programmes are initiated by selecting one or two migratory cells that lead outgrowing sprouts, while a limited number of stalk cells follow these tip cells and account for the actual branch as it forms. Very few cells thus form the initial branches, and later branch growth and extension can be accomplished by further cell divisions (in the vasculature) or by cell growth through polyploidization (in the tracheal system). In contrast to these relatively simple branching systems, branching in other organs such as the kidney or the lung relies on collective behaviour of larger groups of cells (see for example Neumann et al., 2018). At this level of complexity, single cell behaviour is less well understood and, thus far, the branching programme appears to differ from the two systems discussed here. Furthermore, these latter organs generate blind-ended tubes and not interconnected networks, so there are no in-built anastomosis processes.

**Figure 1.**
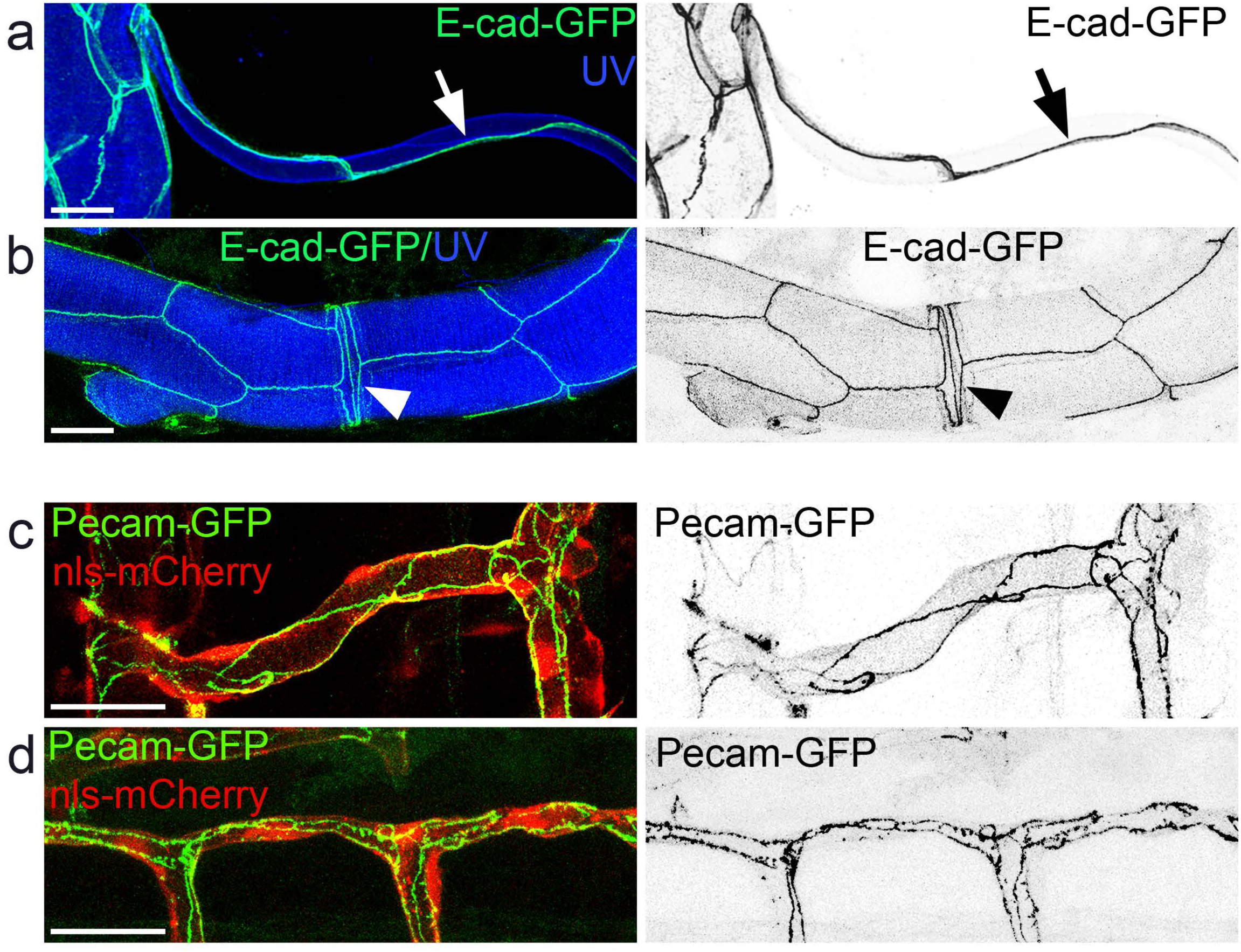
Examples of branched architectures in the Drosophila trachea and the zebrafish vasculature. The architecture of a unicellular tube with autocellular junctions (arrow) of the dorsal branches (a) and doughnut-shaped cells (arrowhead) forming seamless tubes upon tracheal anastomosis (b) are shown in dissected trachea from E-Cad-GFP Drosophila larvae; the lumen is displayed by cuticle autofluorescence in the UV channel. Multicellular tube organisation in intersegmental vessel (c) and dorsal longitudinal anastomotic vessels (d) from 48hpf Tg(fli1a:pecam1-EGFP; fli1a:NLS-mCherry) zebrafish embryos. Scalebars: 30 p.m.

In the following, we will briefly describe the branching process of the developing tracheal system in the drosophila embryo and of the developing vasculature of the zebrafish embryo and the vasculature of different vertebrate organs. We will then discuss similarities as well as differences between the trachea and the vasculature. We will focus on cell behaviour rather than on the molecular control of these branching processes, highlighting the variations in cell behaviours and subsequent tube organisations that ultimately lead to interconnected tubular structures.

## 2 Branching and anastomosis in the Drosophila tracheal system

The insect trachea is a complex branched tubular network, which facilitates oxygen exchange through simple diffusion of air from the outside world to every cell within the organism (Manning and Krasnow, 1993; Samakovlis et al., 1996a). The tracheal system of the Drosophila larva is formed during embryonic development (reviewed in Caussinus et al., 2008; Ghabrial et al., 2003; Lubarsky and Krasnow, 2003; Maruyama and Andrew, 2012; Uv et al., 2003) and stabilized during the final stages via a chitineous apical matrix (Luschnig et al., 2006; Tonning et al., 2005; Wang et al., 2006). Tracheal tubes arise from 10 separate primordial placodes, which form on the surface of each side of the embryo in a segmental fashion. Tracheal cells are determined as groups of epidermal cells and remain fully polarized epithelial cells as the placode invaginates and cells divide one more time (Isaac and Andrew, 1996; Wilk et al., 1996). Subsequently, small subsets of the approximately 80 cells in each placode start to migrate out in a stereotypical manner in different directions, thereby forming distinct branches reaching different organ systems. Specialized fusion cells from adjacent and contralateral segments ultimately contact and establish tubular connections between different metamers or a single metamer, respectively (Samakovlis et al., 1996b). Remarkably, subcellular tubes extend from single terminal cells at the periphery of the branches in a tree-like fashion, enlarging the surface covered by the branched tracheal network and ensuing that every cell in the embryo and the resulting larvae is in contact or in close vicinity to the tracheal system in order to be provided with appropriate levels of oxygen (Guillemin et al., 1996).

## 2.1 Tracheal branch formation through cell migration

Genetic, reverse genetic and live imaging analyses have provided ample evidence that the branching pattern of the tracheal system emerges from and relies upon directed cell migration towards target tissues. Migration of tracheal cells requires the activation of the fibroblast growth factor (FGF) receptor signalling pathway, which is achieved via the ligand Branchless (Bnl) through binding to the FGF receptor Breathless (Btl). While all tracheal cells express Btl (Klambt et al., 1992), only very few, localized non-tracheal cells express the ligand Bnl (Sutherland et al., 1996). Tracheal cells closest to sources of Bnl start to extend filopodia and migrate in the direction of the Bnl source. As they migrate away from the tracheal placode, these so-called “tip cells” pull along “stalk cells”, which results in the formation of distinct tracheal branches. Since all tracheal cells remain fully integrated in the remodelling epithelium during this process, tip cell migration leads to a complete architectural restructuring of the initial epithelial sac, transforming it into a complex, branched epithelial tree. The three-dimensional organisation of the tracheal structures within the developing embryo is due to the spatially restricted Bnl expression and its migration-activating function in tracheal cells via Btl signalling; loss of function mutations in the *btl* or *bnl* genes leads to a complete lack of branching, while ectopic sources of Bnl secretion leads to the formation of ectopic branches towards these sources (Klambt et al., 1992; Sutherland et al., 1996).

Subsequent studies showed that migration of certain branches requires integrin signalling (Boube et al., 2001) as well as Slit/Robo signalling (Englund et al., 2002; LundstrÖm et al., 2004). These findings support the notion that FGF-induced cell migration is at the top of the hierarchy controlling branching via directed cell migration, but that other cues assist FGF signalling either positively or negatively. The highly organized nature of the tracheal branching pattern is also due to topological constraints, e.g. through the restriction of migratory routes by other organs within the embryo (Casanova, 2007).

## 2.2 Tracheal branch elongation through cell intercalation and cell shape changes

Because **cells do not divide** during tracheal branching, elongation of the branch stalk has to be achieved via cell shape changes and/or via cell rearrangements, and these are indeed the two prevailing mechanisms giving rise to the extended shape of functional tracheal branches. Genetic studies combined with live imaging and laser ablation experiments have shown that stalk cells, upon pairing along the proximal-distal axis of the outgrowing branch, start to rearrange and, most strikingly, start to intercalate (Ribeiro et al., 2004; see Fig.2 and Movie 1). During this complex process, each of the paired cells starts to reach around the lumen in-between the next pair of cells and eventually contacts itself at the other side of the tube, initiating the formation of the first junctional self-contacts (also referred to as autocellular junctions). These sites of self-contacts subsequently enlarge as the cells continue to intercalate. At the end of the process, stalk cells have generated large portions of autocellular junctions along the branch axis, while remaining connected to the more proximal and more distal cells through intercellular, ring-link junctional contacts (see Ribeiro et al., 2004 for a more detailed description). For cells being able to intercalate in the described manner, they have to be aligned (paired) along the branch axis at the beginning of the process. The intercalation process is under transcriptional control (Ribeiro et al., 2004) and requires regulation of intracellular trafficking of E-cadherin (Shaye et al., 2008). In addition, two extracellular proteins, Piopio and Dumpy, form a lumenal scaffold which is required to stop the process of self-contact formation at the poles (Jaźwińska et al., 2003). If this process were to be continued to the end, it would result in the complete disruption of branch formation, since individual branch cells would form exclusively self-contacts and would no longer be held together along the branch axis via intercellular connections (Jaźwińska et al., 2003).

**Figure 2.**
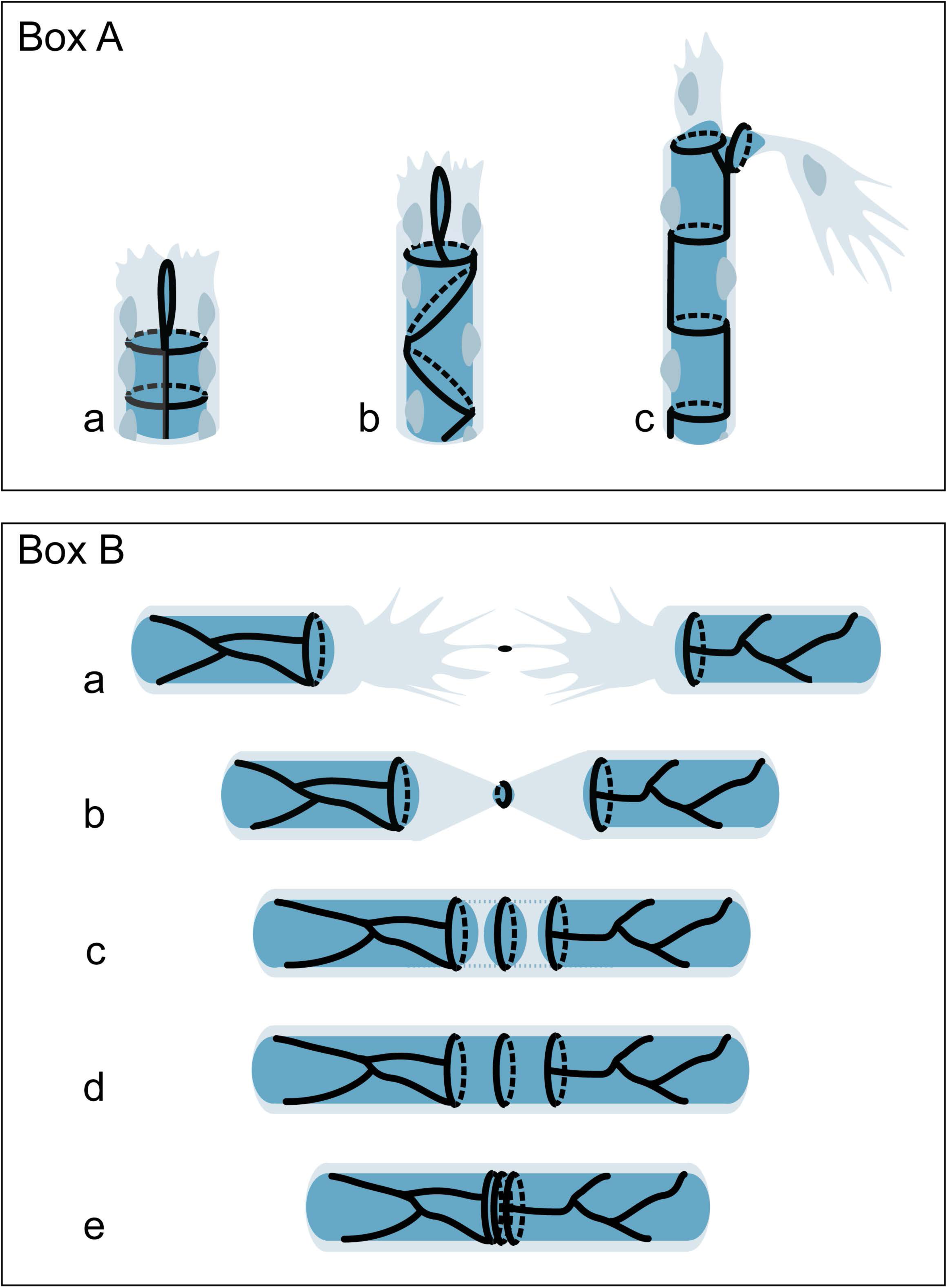
Box A. Branch elongation in the drosophila trachea. Branch extension is driven by active tip cell migration and passive stalk cell intercalation. An initial step requires stalk cell pairing along the proximal-distal axis. In a second step stalks cells start to rearrange and to intercalate. As an outcome, stalk cells have generated autocellular junctions, while they remain connected to their neighbouring cells with intercellular junctional contacts. The cell cytoplasm and nucleus are shown in light blue hues, the lumen in dark blue and cell junctions in black. **Box B. Anastomosis in the drosophila trachea**. Fusion cells of neighbouring branches connect via filopodial contacts (a) upon which a novel apical membrane patch is generated (b), expanded (c) and eventually fuses with the two apical membrane compartments from the proximal side generating a continuous lumen (d-e). The cell cytoplasm is shown in light blue, the lumen in dark blue and cell junctions in black.

How is this striking process of cell intercalation brought about? While many cell intercalation events in the epidermis of the Drosophila embryo rely on junctional shrinking and extension driven by cell autonomous actomyosin activity (Guillot and Lecuit, 2013; Munjal and Lecuit, 2014; Vichas and Zallen, 2011), tracheal stalk cell intercalation is driven by the migrating tip cell in a cell non-autonomous manner (Caussinus et al., 2008). Tip cell migration causes tension on the connected stalk cells due to their attachment to the dorsal trunk; as a result of this tension (or in order to release this tension), tracheal cells intercalate and subsequently extend dramatically until they are organized in a cord-like manner along the branch axis, with the most distal stalk cell remaining attached to the leading tip cell, and the most proximal stalk cell remaining attached to the dorsal trunk. Using a protein degradation method, it has recently been confirmed that actomyosin activity is indeed not required for branch formation in the tracheal system (Ochoa-Espinosa et al., 2017). It thus appears that branch formation is brought about by active tip cell migration and passive stalk cell intercalation, leading to the particular unicellular architecture of most tracheal branches (see Fig.1a and Fig.2c).

## 2.3 Making connections between tracheal branches

The two distal cells of most tracheal branches subsequently differentiate into a terminal and a fusion cell, respectively, a process that is driven by external signals and results in these two specific and distinct cell types (Caviglia and Luschnig, 2013; Chihara and Hayashi, 2000; Samakovlis et al., 1996b). The fusion cell is responsible to connect neighbouring metameres (or the left and right side of the embryo within a metamer), thereby generating an interconnected luminal space (Beitel and Krasnow, 2000; Samakovlis et al., 1996b reviewed in Caviglia and Luschnig, 2014; see Fig.3). Fusion cells of neighbouring branches connect via filopodial contacts, which result in the deposition of E-cadherin and the establishment of a novel cell-cell contact site and the subsequent insertion of a novel apical membrane patch (Lee, 2003; Lee and Kolodziej, 2002a). E-cadherin recruits several cytoskeletal proteins and triggers the formation of an actin- and microtubule-rich filamentous structure, which connects the two apical surfaces of each fusion cell (Tanaka et al., 2004). This structure serves as a scaffold for vesicle transport and helps to assemble or enlarge the apical membranes (Jiang and Crews, 2007; Kato et al., 2016). The two apical membrane compartments eventually need to fuse in order to generate a continuous lumen, and it has been shown that this requires a specialized endosomal/lysosomal related organelle (LRO; Caviglia et al., 2016). Once this has happened in both fusion cells, a continuous lumen connects the adjacent or contralateral metamers. Fusion cells are transformed during this process into hollow cylinder-like cells, making ring-like adherens junctional contacts with the two neighbouring tracheal cells. Since the two junctional rings are brought into close vicinity at the end of the anastomosis process and border an apical membrane spanning the entire luminal circumference without a seam, these cells have been described as “doughnut” cells (Uv et al., 2003), and they remain embedded in this particular configuration within the tracheal system during all subsequent larval stages (see Fig.3).

**Figure 3.**
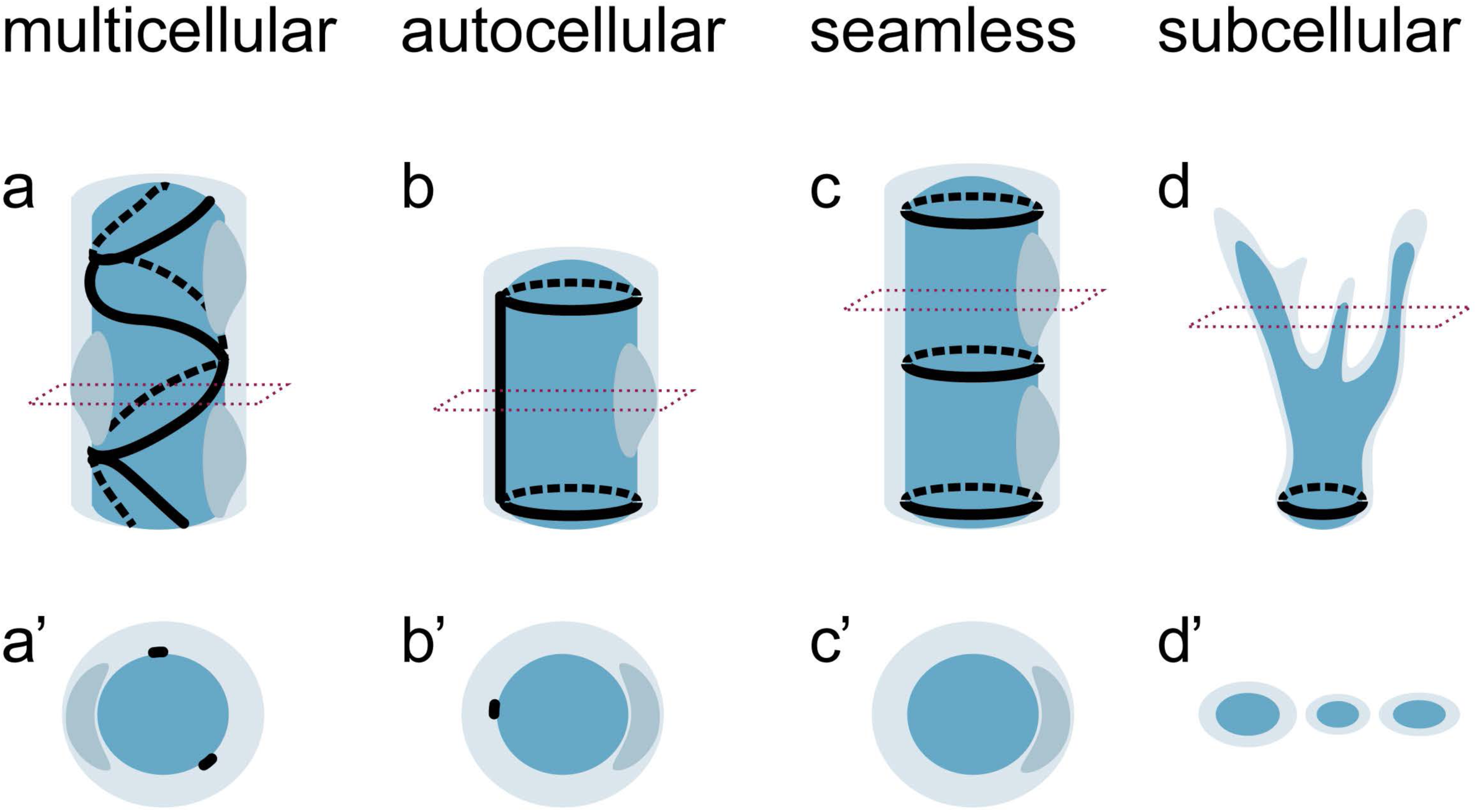
Tube architectures in the trachea. Depictions of variations of tube organisation: multicellular (a), autocellular/unicellular (b), seamless (c) and subcellular (d), in side view (a-d) and transverse view (a’-d’). The cell cytoplasm and nucleus are shown in light blue hues, the lumen in dark blue and cell junctions in black. Red dotted lines display the cross-sections for a’-d’.

## 2.4 Tube architectures in tracheal branches

As described in the previous sections, the branching process in the drosophila embryonic trachea transforms an epithelial sac into a complex tubular network built by four types of tubes displaying different architecture (Uv et al., 2003) (see Fig.4). The main airway structure, the dorsal trunk, which extends throughout the embryo along the anterior-posterior axis, is made up from the largest part of the cell pool of each tracheal placode and is characterized by its ***multicellular tube*** architecture (Fig.4a). Most metameric branches are made up of cells that seal the luminal space with autocellular junctions (self-contacts), a configuration that results from the intercalation process during branch elongation described above. Such tubes have been described as ***autocellular tubes*** (Fig.4b). During the process of tracheal anastomosis, doughnut-shaped cells are formed which wrap around the lumen and seal it without adherens junctions, thereby forming a ***transcellular tube*** (also referred to as a ***seamless tube)*** (Fig.4c). At the periphery of the tracheal system, additional fine tubes are formed within numerous and highly ramified cytoplasmic extensions each carrying an internal tube, blunt-ended at the tip of the extension but connected in the body of the terminal cell with the apical membrane and the luminal space. These terminal cells therefore contain a large tree of extensions containing a ***subcellular tube*** (Fig.4d).

**Figure 4.**
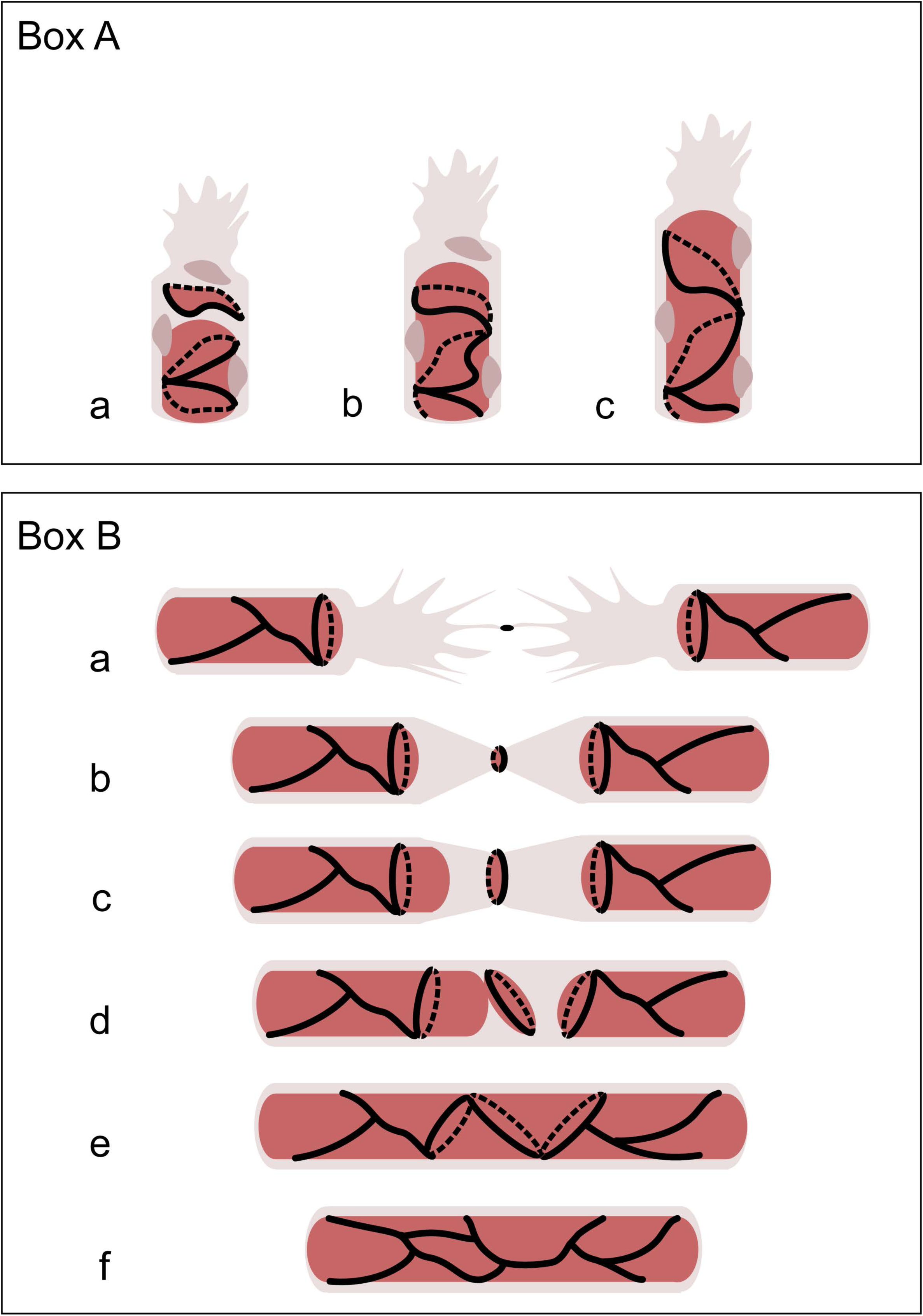
Box A. Branch elongation in zebrafish vasculature. In a first step, stalk cells follow the migrating tip cells in a chain-like organisation and connect to each other via adherens junctions. During sprouting, stalk cells elongate and expand their junctional contacts with neighbouring cells. As a result, stalk cells of the sprout appear in a multicellular organisation. The cytoplasm is shown in light pink, the lumen in dark pink and cell junctions in black. **Box B. Anastomosis in the zebrafish vasculature**. Two tip cells generate an initial site of cell junctions upon filopodial contact, (a), where apical membrane appears (b) and further expands into a disc-like structure (c). During anastomosis, membrane invagination generates a transcellular lumen (d) which eventually fuses with the *de novo* generated distal apical membrane compartment at the contact site, thereby forming a transient unicellular tube within the anastomosing cell (e). Subsequently, cell rearrangements lead to the formation of a multicellular tube (f). During anastomosis via cord hollowing, the membrane compartments are assembled by cell rearrangements and junctional remodelling, thereby generating a multicellular tube.

## 3 Branching and anastomosis in the early zebrafish vasculature

While the tracheal system of insects transports gases throughout the embryo and this transport relies to a large extent on passive diffusion, the vasculature of vertebrates is a transport conduit for cells and fluids and this transport is assisted by a pump, the heart. During a process called vasculogenesis, an initial simple network of tubes is connected to the beating heart via the assembly of endothelial cells and the formation of tubular structures (Herbert and Stainier, 2011; Schuermann et al., 2014; Szymborska and Gerhardt, 2017). Most of the additional vascular branches are subsequently added by the process of angiogenesis, the sprouting of novel vessels from pre-existing vessels (Betz et al., 2016; Hogan and Schulte-Merker, 2017; Schuermann et al., 2014; Szymborska and Gerhardt, 2017); angiogenesis can thus be considered as a process of branching morphogenesis.

Although the vasculature of larger vertebrate embryos consists of millions of endothelial cells, it turns out that the individual sprouting and anastomosis events during the process of angiogenesis are brought about by a limited number of cells, often by either one or two tip cells (sprouting) and by two fusion cells (anastomosis) and appears to be similar to the corresponding processes in the tracheal system of insects. Careful analyses of endothelial cells in fixed vascular beds in combination with live imaging studies have provided much insight into the cellular events occurring during angiogenesis. We will briefly describe these findings, and then compare branching morphogenesis in the trachea and the vasculature.

## 3.1 Vascular sprout formation through cell migration

In a landmark paper published in 2003 (Gerhardt et al., 2003), it was shown that Vascular Endothelial Growth Factor A (VEGF-A) controls angiogenic sprouting in the early postnatal retina in mice by guiding the migration of specialized endothelial cells at the tip of vascular sprouts through the formation of filopodial extensions. While tip cells respond to VEGF-A by guided migration as a result of a gradient of VEGF-A, stalk cells respond by increased proliferation triggered by a concentration threshold of VEGF-A. In this scenario, branch direction and stalk extension are coordinated by different agonistic responses to the same ligand (Gerhardt et al., 2003). In more recent years, several other guidance cues acting on endothelial tip cells have been identified, in particular cues that are also involved in axonal guidance (Eichmann and Thomas, 2013; Eichmann et al., 2005; Lu et al., 2004; Torres-Vázquez et al., 2004), adding to the general picture that the sprouting process is to a large extent guided and controlled by directed cell migration.

## 3.2 Vascular sprout elongation through cell recruitment, cell division and cell rearrangement

As already highlighted in the study of Gerhardt and colleagues (Gerhardt et al., 2003), sprout extension is in part brought about by the division of stalk cells. In other cases, such as during the formation of segmental arteries in the zebrafish embryo, tip cells also divide at rates equal to stalk cells. During the early events of sprout formation in the segmental arteries, stalk cells are also recruited from the parental vessel, the dorsal aorta; they are drawn into the stalk in a chain-like organisation as they follow the migrating tip cells to which they are connected via adherens junctions (Hogan and Schulte-Merker, 2017). The combination of cell recruitment and proliferation thus leads to segmental sprouts that on average contain four endothelial cells (Nicoli et al., 2012). Furthermore, asymmetric endothelial cell divisions have been shown to enable migration of the angiogenic sprouts via differential VEGF receptor activity (Costa et al., 2016). Therefore, and in sharp contrast to tracheal branch elongation, cell division and the resulting increase in cell number contributes to a large extent to the branch elongation process in the developing vasculature.

In addition to cell proliferation and migration, the sprouting of segmental arteries is also associated with cell elongation, which changes the cellular configuration from a serial to a parallel arrangement. These cell rearrangements lead to an extensive longitudinal pairing, particularly of the stalk cells, which allows them to form an intercellular luminal space. Ultimately, this process leads to the formation of a multicellular tube, where a cord hollowing process forms the lumen (see Fig.5).

**Figure 5.**
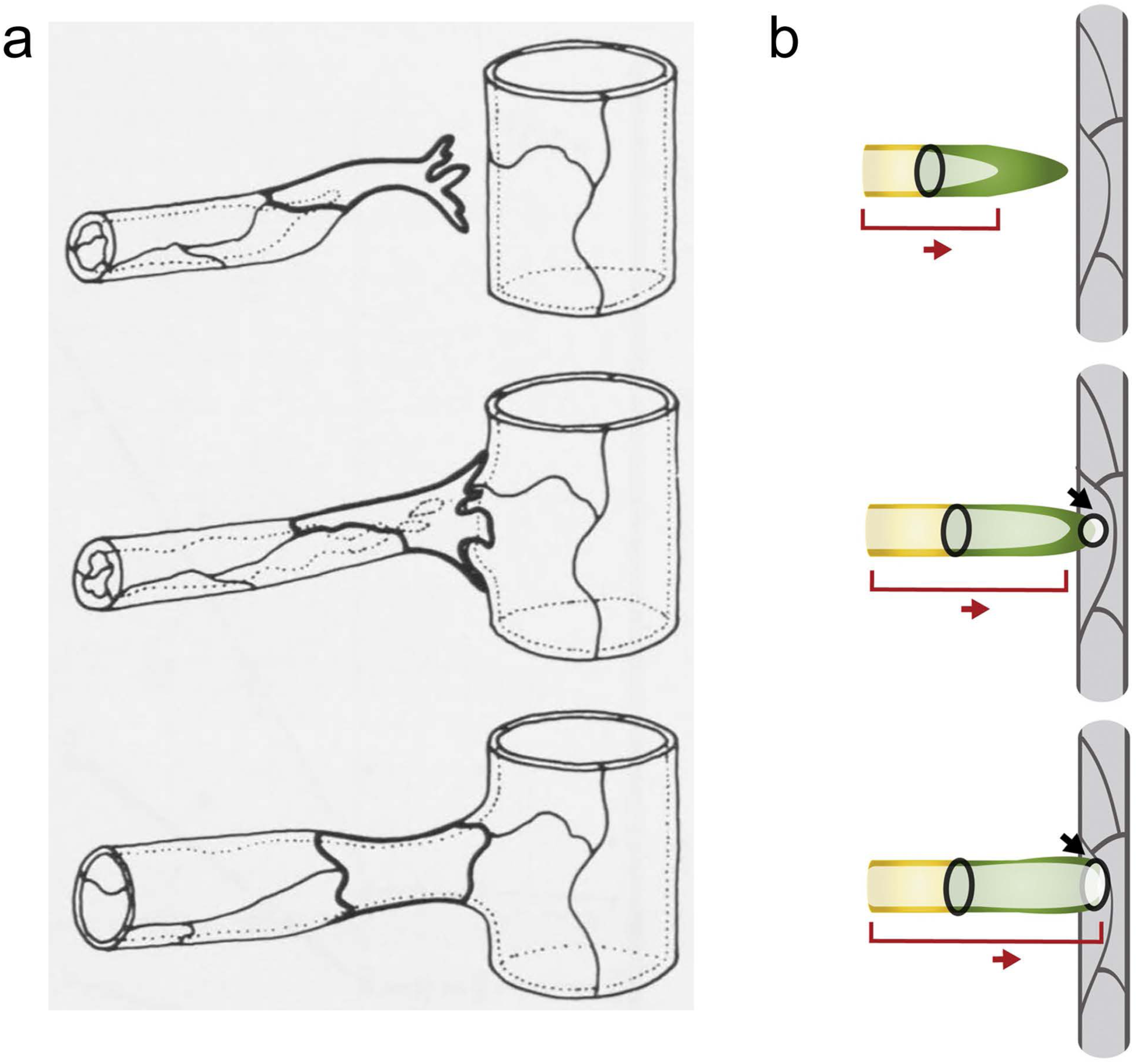
Seamless endothelia in electron microscopy studies. Diagram showing a growing sprout from a capillary making contact with a venous vessel generating a seamless tube (a). Picture taken from Wolff and Bär, 1972. A model of vessel fusion in the embryonic zebrafish brain (b). A tip cell (green) which is followed by stalk cells (yellow) connects (black arrow) to a functional vessel (grey) and generates temporarily a seamless tube. Lumen is shown in bright green, bright yellow, red bar. Picture taken from Lenard et al., 2013.

Ablation of VE-cadherin/Cdh5 or interfering with F-actin polymerization inhibits stalk cell elongation and consequently prevents multicellular tube formation (Sauteur et al., 2014). In contrast to Drosophila tracheal cells, stalk cell elongation appears to be an active process. Consistent with this view, ablation of the tip cell in the nascent sprout does not lead to a detectable stalk retraction (Sauteur et al., 2014). Closer analyses of junctional and F-actin dynamics have uncovered a novel mechanism by which endothelial cells can use each other as substrates for cell migration and elongation (Paatero et al., 2017). The cells use oriented, oscillating lamellipodia-like protrusions – so-called junction-based lamellipodia or JBL, which emanate from and are connected to adherens junctions. High-resolution in vivo imaging and genetic analyses showed that JBL provide an oscillating ratchet-like mechanism, which is used by endothelial cells to move along or over each other and thus provides the physical means for cell rearrangements. By immunofluorescence, junctional protrusions that may be related to JBLs, have also been described in the postnatal mouse retina (Cao et al., 2017), raising the possibility that JBLs may provide a general mechanism of endothelial cell rearrangements.

## 3.3 Making connections between vascular sprouts

The amenability of the early zebrafish embryo for high-resolution live imaging makes this system particularly attractive for a characterization of the cellular activities involved in vascular anastomosis. Interestingly, it was found that sprout connection and lumenisation occurs via two distinct mechanisms, transcellular lumen formation and cord hollowing (Herwig et al., 2011). During the initial steps of anastomosis, newly formed endothelial cell junctions at the initial site of tip cell filopodial contacts expand into ring-like structures, generating an apical membrane pocket between the two sprouts. During the *cord hollowing* process, which occurs mostly when two non-lumenized sprouts connect, these apical membrane compartments are brought together via cell rearrangements accompanied by extensive junctional remodelling, which eventually leads to lumen coalescence and the formation of a multicellular tube. Vessel anastomosis via *transcellular lumen formation* occurs in lumenized sprouts with blood pressure being the major driving force which leads to membrane extension/invagination into the tip/fusion cell, thereby generating a transcellular lumen (Gebala et al., 2016). This transcellular lumen eventually fuses with the de novo generated distal apical membrane compartment at the contact site, thereby forming a transient unicellular tube in the anastomosing cell. This transcellular tube segment is, however, rapidly transformed into a multicellular tube via cell rearrangements and cell splitting (Lenard et al., 2013; see Fig.6). As a result, both mechanisms of vascular anastomosis, cord hollowing and transcellular lumen formation, lead to the formation of multicellular tubes, and no apparent sign remain as a testimony that a particular segment of a given branch has been formed via the anastomosis of two individual sprouts.

**Figure 6.**
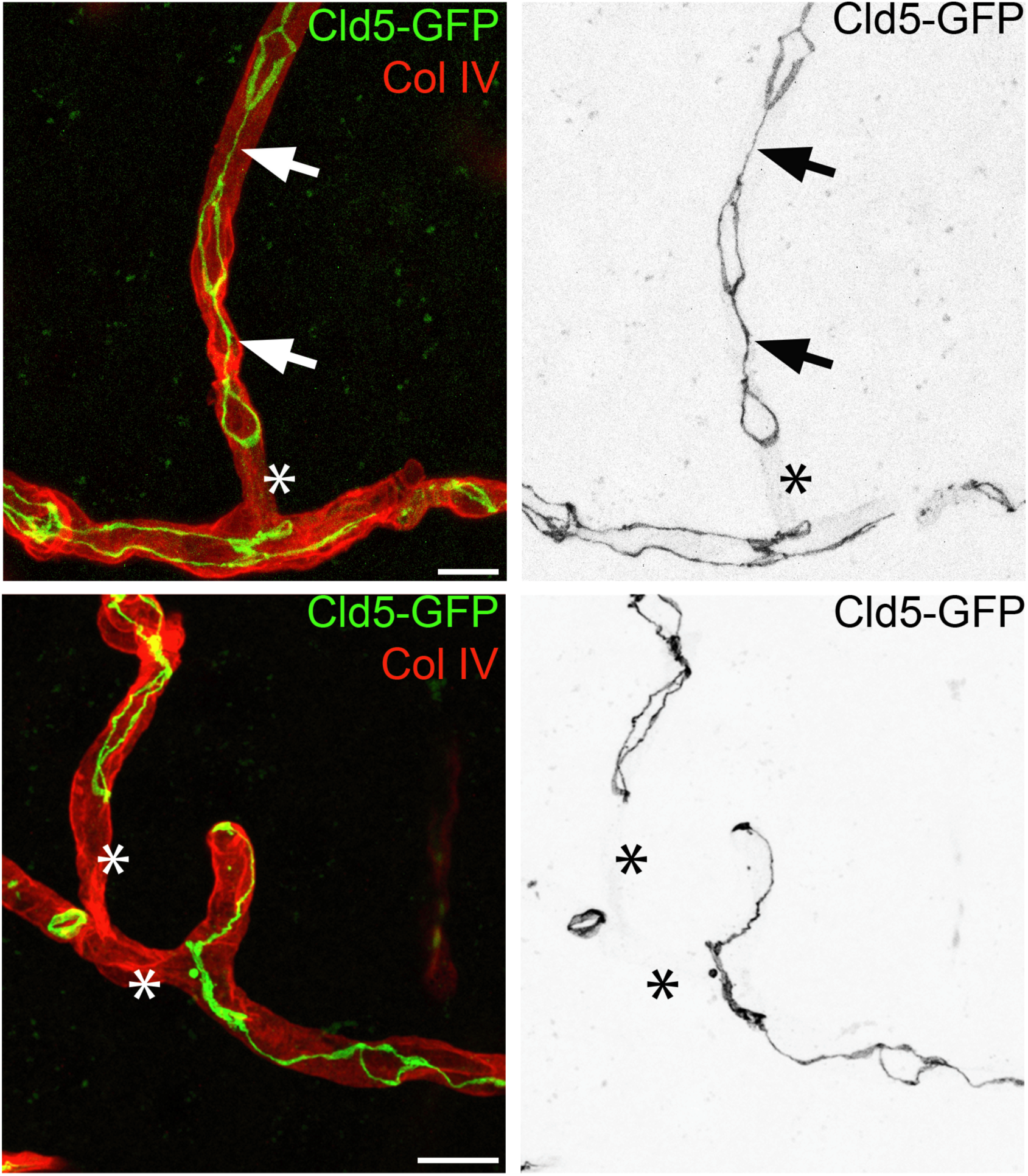
Seamless tubes and tubes with autocellular junctions in light microscopy studies. Multicellular, autocellular junctions (arrows) and seamless stretches (asterisks) in brain capillaries of a 3 months old Tg eGFP-Claudin5 (Knowland et al., 2014) (green) mouse in C57Bl6/J background. The endothelial basement membrane is visualized with Collagen IV staining (red). Scalebars: 10 p.m.

## 3.4 Tube architectures in vascular branches

As described above, the common theme of the architecture of vascular tubes is their multicellular nature, both in the developing retina in mice and in the developing vasculature in the zebrafish embryo, two of the most widely used systems to study angiogenesis (Franco et al., 2015; Lenard et al., 2013). This is in sharp contrast to what is seen in tracheal sprouts in the early Drosophila embryo, where most branches are formed by autocellular, transcellular (seamless) or subcellular tubes. During angiogenesis, transcellular (seamless) tubes are also formed during cell division in elongating sprouts (Aydogan et al., 2015) and during membrane invagination in anastomosing vessels (Lenard et al., 2013); however, these tubes are transient and remodel to become multicellular tubes. Subcellular tubes in the trachea are blunt-ended, and such tubes most likely do not exist in the vasculature as they cannot support blood flow and would undergo pruning.

## 4 Similarities and differences in cell activities underlying the formation of two distinct branched organs

## 4.1 Directed cell migration

As outlined above, the complex branching pattern of both the Drosophila tracheal system and the developing vasculature (as formed via angiogenesis) is to a large extent controlled by directed cell migration. In both cases, so-called tip cells at the periphery of the outgrowing branch navigate under the control of ligands secreted by surrounding tissues, mainly ligands of the FGF family in the trachea and the VEGF family in the vasculature (Gerhardt et al., 2003; Klambt et al., 1992; Sutherland et al., 1996), respectively, which are assisted by attractive and repulsive signals also known for their role in neural guidance (Eichmann and Thomas, 2013; Englund et al., 2002; Lundström et al., 2004; Torres-Vázquez et al., 2004). Strikingly, Notch/Dll signalling has been involved in tip/stalk cell selection both in the trachea and in the vasculature (Ghabrial and Krasnow, 2006; Siekmann and Lawson, 2007; reviewed in Phng and Gerhardt, 2009); symmetry breaking is thus achieved by very similar mechanisms in the two systems.

## 4.2 Branch elongation

For branches to connect and fuse with other branches, they have to extend in length and during this process, there appear to be considerable differences in the developing trachea when compared to the zebrafish vasculature. In tracheal branches, stalk elongation occurs in the absence of cell division and is brought about by cell intercalation and cell elongation, both triggered by the tension force exerted onto the stalk cells by the migrating tip cell (Caussinus et al., 2008). Intercalation is passive, meaning that stalk cells do not regionally deploy actomyosin forces to remodel their junctions as a driving force for intercalation (Ochoa-spinosa et al., 2017). As a result of the intercalation process and of the small number of cells participating in it, most tracheal branches present a specific cellular architecture, being made up of cells aligned along the axis of extension in a chain-like manner, sealing the lumen via autocellular junctions and connecting to proximal and distal neighbours via ring-shaped intercellular junctions (see Fig.1,2&4).

In sharp contrast to tracheal branches, vascular sprouts have been shown to form multicellular tubes and their extension is assisted by division of stalk cells and by active cell elongation. Both actin and actomyosin are required for these processes (Sauteur et al., 2014; Phng et al., 2015; Paatero et al., 2017). The tension exerted by the migrating tip cell is thus not interpreted in the same way as in the tracheal system; endothelial cells might react to the tension generated by tip cell migration via increased cell division and enhanced cell elongation rather than via cell intercalation and the generation of autocellular junctions. Consequently, the architecture of the resulting branch is quite different between these two branched organs.

## 4.3 Branch connection

The anastomosis of contacting sprouts involves filopodial contacts between two tip cells as an initial trigger, which leads to the formation of novel cell-cell junctions at the contact site and the formation of a novel apical membrane patch. In the trachea, the connection of the two branches is established via specialized tip cells, the fusion cells, which differ in the expression of molecular markers from the other tip cell fate, the terminal cells (Caviglia and Luschnig, 2013; Chihara and Hayashi, 2000; Samakovlis et al., 1996b). In contrast, the tip cells driving the anastomosis process in the vasculature appear to be more flexible, exchange in certain cases position with stalk cells, and so far, no distinction has been made between tip cell fates during branch elongation and during anastomosis. Nonetheless, the connection of the two branches strictly requires cell adhesion molecular markers, and more specifically, E-cadherin in the tracheal system (Lee, 2003) and VE-cadherin and Endothelial cell-selective adhesion molecule a (Esama) in the vasculature. VE-cadherin and Esama are required in a somewhat redundant fashion, to initiate proper contact formation and apical membrane insertion (Sauteur et al., 2017). Despite the differences in the specification of the tip cell fate between the two systems, the common theme during branch connection is filopodial contacts mediated by cell adhesion molecules.

## 4.4 Lumen formation

Much has been learned about the formation of an interconnected apical luminal space in tracheal fusion cells from studying the developing tracheal system (summarized in Caviglia and Luschnig, 2014; Sundaram and Cohen, 2017). Briefly, filopodial contacts between fusion cells leads to the focal accumulation of E-cadherin. Through the insertion of apical polarity complexes and apical proteins and membrane, the spot-like E-cadherin expands into a ring-shaped adherens junction between the two fusion cells, resulting in bipolar epithelial cells. The two apical domains are connected by an F-actin track and microtubule (MT) bundles, requiring the expression of Short stop (Shot), a large, bifunctional protein containing actin- and MT-binding domains (Lee and Kolodziej, 2002b). This central track may provide a highway for transporting vesicles and organelles by MT-dependent motor proteins. Moreover, an elegant recent study showed that the exocytosis of a lysosome-related organelle (LRO) is involved in the fusion of the two apical membrane compartments within the fusion cells (Caviglia et al., 2017).

Little is known at the molecular level about how the apical luminal membrane is generated in the developing vasculature during angiogenesis in vivo. It has been proposed that the lumen arises from the fusion of intracellular vacuoles, which eventually fuse in-between adjacent cells (Kamei et al., 2006; Yu et al., 2015). Another study instead proposed that hemodynamic forces dynamically shape the apical membrane to form and expand new lumenized vascular tubes (Gebala et al., 2016; Lenard et al., 2013). Future studies will have to address which cellular compartments contribute membrane to the growing apical domain and which trafficking pathways are involved in this phenomenon and how they are regulated.

## 5 A look back and a look ahead; even more similarities?

Studies of the Drosophila tracheal system revealed that the epithelial tubes formed during the branching process are made up of four different types of tubes, differing in the architectural arrangement of the individual constituting epithelial cells (Fig.4a to d). Similar studies on the developing vasculature in the zebrafish embryo and in the developing mouse retina showed that most of the vascular tubes are multicellular tubes in these systems. Using live imaging, cell intercalation leading to stable autocellular junction formation has not been observed thus far in the vasculature. Strikingly, bipolar cells similar to the seamless doughnut cells in the drosophila trachea are transiently formed during vascular anastomosis, but their architecture is remodelled as cell rearrangements followed by cell fission events convert the seamless tube into a multicellular tube. Tip cell division can also lead to doughnut-like cells, but cell rearrangements also convert this architecture into a multicellular tube (Aydogan et al., 2015). Stable doughnut-like cells have not been seen yet in the developing vasculature in the developing zebrafish embryo or in the retina.

However, most of the vascular beds that have been analysed with regard to their cellular architecture in live embryos represent structures that continue to remodel and in which endothelial cells are still actively dividing. What about the capillary networks in fully developed, adult organs? Using light and electron microscopy, it has been reported many years ago that capillaries in these adult vascular beds are made up to a large extent by seamless tubes (Wolff, 1964). It has been estimated that as many as 30-50% of the tubes in brain capillary beds are represented by seamless tubes (Wolff and Bär, 1972; reviewed in Bär et al., 1984; see also Fig.7), and such tubes have also been found in many organs and tissues studied (different types of muscle, brain, duodenal villi, renal glomeruli, but not in normal liver, spleen and endocrine glands). Furthermore, it has also been reported that many endothelial cells form “a cuff around the capillary lumen and make contact with themselves along the longitudinal axis of the vessel” (Bär et al., 1984). Such stretches of tubes would correspond in their architecture to those seen in the developing trachea and referred to as autocellular tubes therein (see Fig.3). Based on the positioning of seamless tubes within the network, it has been suggested that seamless tubes develop via the anastomosis of venous cells with pre-existing vessels (Bär et al., 1984; Fig.7). Furthermore, some ideas about how seamless tubes and tubes with autocellular junctions might arise have already been discussed in the literature (see Strilić et al., 2010).

More recently, light microscopy analyses in the adult mouse brain and retinal vasculature using fluorescent junctional markers have indeed shown the presence of autocellular junctions and seamless tubes (see examples in Fig.8). Generally, these observations suggest that while all arteries and veins are built as multicellular tubes, the microvasculature (arterioles, venules and capillaries) show both multi- and unicellular composition, the latter with autocellular junctions, as well as interspersed stretches of seamless tubes (MA-M and CB, unpublished data).

These findings raise a number of interesting questions. Do autocellular junctions in the adult vasculature form in a manner similar to how they form in the developing tracheal system, i.e. through tension-releasing cell intercalation? If yes, are protein scaffolds similar to the ones formed by Pio and Dumpy in the trachea also required to halt intercalation? And do stable seamless tubes arise via anastomosis and/or cell division, as has already been described in vessels that subsequently remodel and become multicellular? Further studies describing the formation of such terminal capillary beds using live imaging analyses should provide insight into how such autocellular and seamless tubes form. It is possible that these studies will reveal that there are even more similarities between the cell behaviour of the Drosophila tracheal system and the vasculature in mammals than we already see. Whether such similarities in cell behaviour and tube architecture will also be accompanied by similarities in molecular regulation remains to be seen, as we learn more about each of these fascinating structures. It also remains to be seen whether the identification of molecular players involved in the formation of terminal capillary beds might reveal new approaches for treating vascular diseases, such as ischemia, or might allow to develop novel strategies to increase or decrease capillary density in organ systems in health and disease.

## Acknowledgments

Work from the MA laboratory was supported by the Swiss National Science Foundation and by the Kantons Basel-Stadt and Basel-Land. Work from the CB laboratory was supported by the Swedish Cancer Society (grant numbers CAN-2016-0777, 150735); the Swedish Research Council (grant number 2015-00550]; the Leducq Foundation [grant number 14CVD02); and the Knut and Alice Wallenberg Foundation (Grant number 2015.0030).

## Movie 1

Animation of the cell intercalation process in the drosophila dorsal branches, which leads to an extended tube with autocellular junctions. A red and a blue cell are originally in a paired configuration. Both cells start to reach around the lumen, but on opposite sides along the tube axis. Once either the red or the green cell reaches around the lumen, it touches itself and established autocellular contacts (also called self-contacts). As the cells intercalate further, these autocellular junctions elongate and replace the intercellular junction, until each cell seals the lumen with autocellular contacts and remains connected to the neighbouring distal and proximal cells with ring-like intercellular junctions. See Ribeiro et al., 2004 for more details.

